# The invasive soft coral *Xenia umbellata* has been confirmed in Cuban waters

**DOI:** 10.64898/2026.06.08.730793

**Authors:** Nikolaos V. Schizas, Daniel A. Toledo-Rodriguez, Nilda M. Jimenez Marrero, Alex J. Veglia, José Espinosa Sáez, Reinaldo Estrada Estrada, Amarilys Martínez Pérez, Yamile del Carmen Luguera González, J. David Muñoz-Maravilla, Ernesto Weil, Catherine S. McFadden

## Abstract

Octocoral colonies with unusual morphology were detected in September 2022 and October 2023 in two coastal areas east of Havana, Cuba, and tentatively identified as *Unomia stolonifera. U. stolonifera* is an invasive octocoral from the Indo-Pacific that was first reported in the Caribbean off Venezuela in the 2000’s, where it has spread rapidly, smothering coral reefs and substantially altering benthic communities. After obtaining tissue samples from a Cuban octocoral colony, we re-examined the specimen using molecular barcoding of three mitochondrial regions (16S/ND2, mtMutS, COI) and the nuclear large ribosomal subunit (28S rRNA), and we unequivocally identified it as *Xenia umbellata. X. umbellata*, a native of the Red Sea, was first identified in southern Puerto Rico in October 2023 and has since been found in various marine ecosystems along the southern coast of the island. The presence of either invasive octocoral species in Cuba or elsewhere in the Caribbean would be of a serious environmental concern due to their documented tolerance, totipotentiality, propagation capacity and significant negative interactions with local benthic fauna. Attempts to eradicate the invasive soft coral colonies from Cuban waters have been initiated with apparent success, helping to control further expansion. The most likely introduction pathway is the accidental or intentional releases from the aquarium trade but transport via ballast water cannot be ruled out. We cannot discount the possibility of independent invasion events from different routes to Puerto Rico and Cuba occurring within a year of each other. Propagation from Cuba to Puerto Rico, or vice versa, which we consider highly improbable, would likely imply that soft coral populations may also have been established on Hispaniola but have remained undetected in the Dominican Republic and Haiti to date.

## Introduction

In September 2022, during an ecological assessment for the ongoing project “Habana Sumergida” (Underwater Havana) conducted by the Center of Marine Research at the University of Havana (CIM-UH), several colonies of an atypical crustose octocoral were encountered near Havana, Cuba. These sightings occurred in the Bacuranao Inlet, Havana del Este municipality (82°14’39.7” W; 23°10’47.4” N) (Fig. 1) with an approximate area of 1 m^2^ and at a depth between 8 and 11 meters. The stolons were attached to the rocky matrix, encrusted with a layer of calcareous red algae. Samples of the unidentified octocoral species were collected and preserved by the Institute of Marine Sciences (ICIMAR) in 4% formalin and deposited in the collections of the National Aquarium of Cuba and the Center for Marine Research at the University of Havana (CIM-UH) with an additional sample preserved in 90 % ethanol for molecular identification. Based on gross morphological characteristics, the octocoral specimens appeared morphologically similar to *Unomia stolonifera* (Gohar, 1938) (Malacalcyonacea: Xeniidae), an invasive soft coral species previously reported from Venezuela (Ruiz-Allais *et al*. 2014, 2021; Espinosa *et al*. 2023).

**Figure 1.**
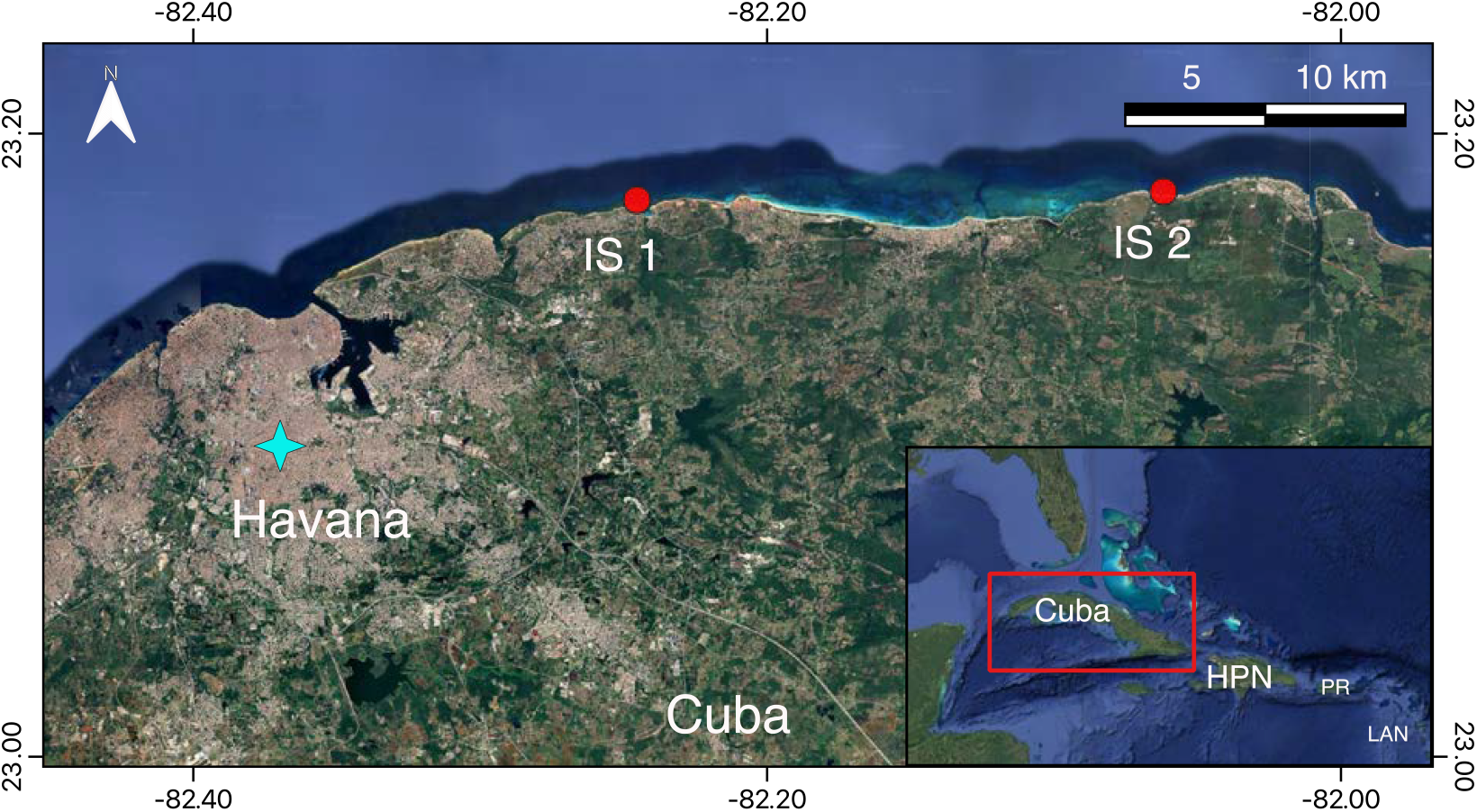
Map showing the geographic location of established colonies of *Xenia umbellata* along the northwestern coast of Cuba. Location of two invasion sites: Bacuranao (IS 1) and Punta Calderas (IS 2). Inset showing the location of Cuba within the Greater Caribbean.

Given the previously reported damage caused by recent octocoral invasions in the Caribbean like that of *Unomia stolonifera* in Venezuela (Ruiz-Allais *et al*. 2014, 2021), to protect Cuba’s coral reef biodiversity eradication efforts were soon after initiated using knives and nylon bags (Espinosa *et al*. 2023; under permits granted by Oficina de Regulación y Seguridad Ambiental, Havana). Despite the eradication of the first colonies, subsequent reef surveys carried out in the Bacuranao Inlet (on February 17 and 23, and March 13, 2023), detected the regrowth and expansion of the invasive octocoral patch near the original sighting location. The reestablished patch overgrew a small portion of the western reef of the channel, on the wall and surface of a surge channel. Although there have been no recent visits by members of CIM-UH to the area, local underwater fishermen and divers report that they have not seen any more colonies.

Expanding the search area, on October 5, 2023, based on the observations of a recreational diver (Mitchell Morales), a team of specialists and divers from the Institute of Marine Sciences (ICIMAR), the Antonio Núñez Jiménez Foundation for Nature and Humanity (FANJ), the Center for Marine Research (CIM-UH), the Rincón de Guanabo Protected Natural Landscape Protected Area (PNPRdG), and associated divers surveyed Boca de Calderas, Mayabeque (82° 3’ 39.21” W; 23° 10’ 57.14” N, Fig. 1). These surveys revealed a second aggregate of octocorals (Figs. 2, 3, 4, 5) now to the east of the city of Havana resembling morphologically the specimens that were reported in Bacuranao Inlet (Figs 3, 4, 5). This new site is located approximately 19 km from the previously reported pulse coral patch in Bacuranao (see Fig. 1), which had been apparently eradicated. The Boca de Calderas aggregate was larger than the one at Bacuranao, covering an estimated area of 150 m^2^. The population extended ∼30 m along the edge of the first submarine terrace at depths of 8 – 13 m. Within this area, 3 to 5 soft coral aggregations were observed, each occupying a surface area of 5 to 6 m^2^. The main aggregation covered about 4 m^2^ and extended over rocks and corals (Fig. 2). The coverage percentage of the invasion was normally above 95% of the occupied area. The colonies in this locality, treated with chlorine tablets covered with nylon, have not been completely eradicated, but so far remain under control with approximately the same distribution.

**Figure 2.**
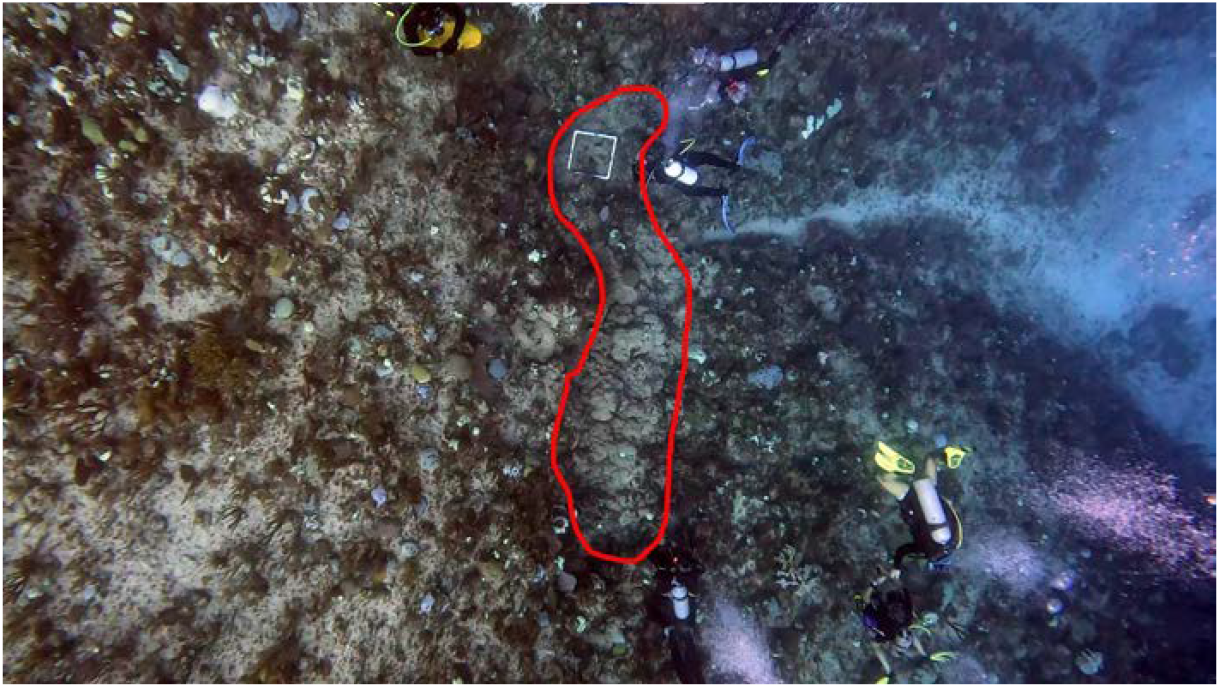
Approximate outline (in red) of the main octocoral aggregate found in Boca de Calderas, Mayabeque, October 7, 2023. Several divers can be seen surveying the site. As reference, the size of the white quadrant is 50 × 50 cm. Photo credit: R. Espinosa Sáez, J. Estrada Estrada, A. Martínez Pérez, Y. del Carmen Luguera González.

**Figure 3.**
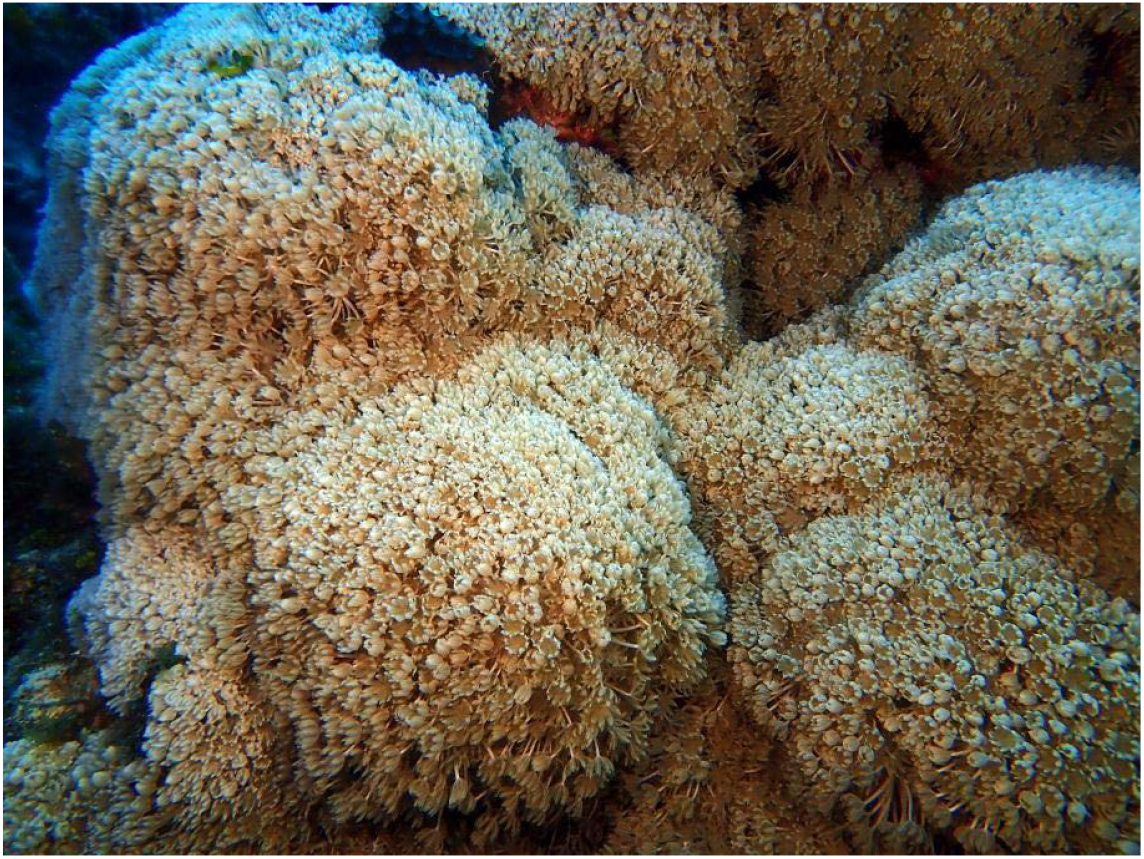
Partial view of an aggregate of *Xenia umbellata* in Boca de Calderas, Mayabeque, October 5, 2023. Photo credit: R. Espinosa Sáez, J. Estrada Estrada, A. Martínez Pérez, Y. del Carmen Luguera González.

**Figure 4.**
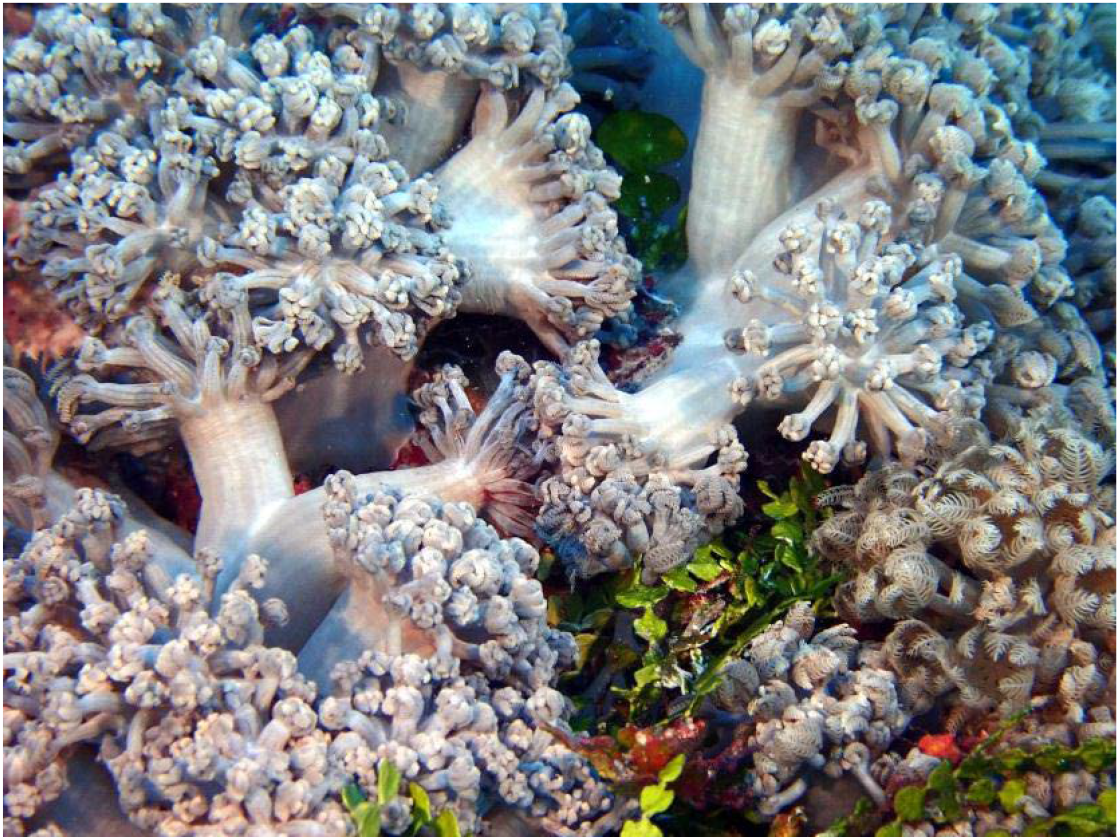
Details of colonies *of Xenia umbellata* showing contracted polyps in Boca de Calderas, Mayabeque, October 5, 2023. Photo credit: R. Espinosa Sáez, J. Estrada Estrada, A. Martínez Pérez, Y. del Carmen Luguera González.

**Figure 5.**
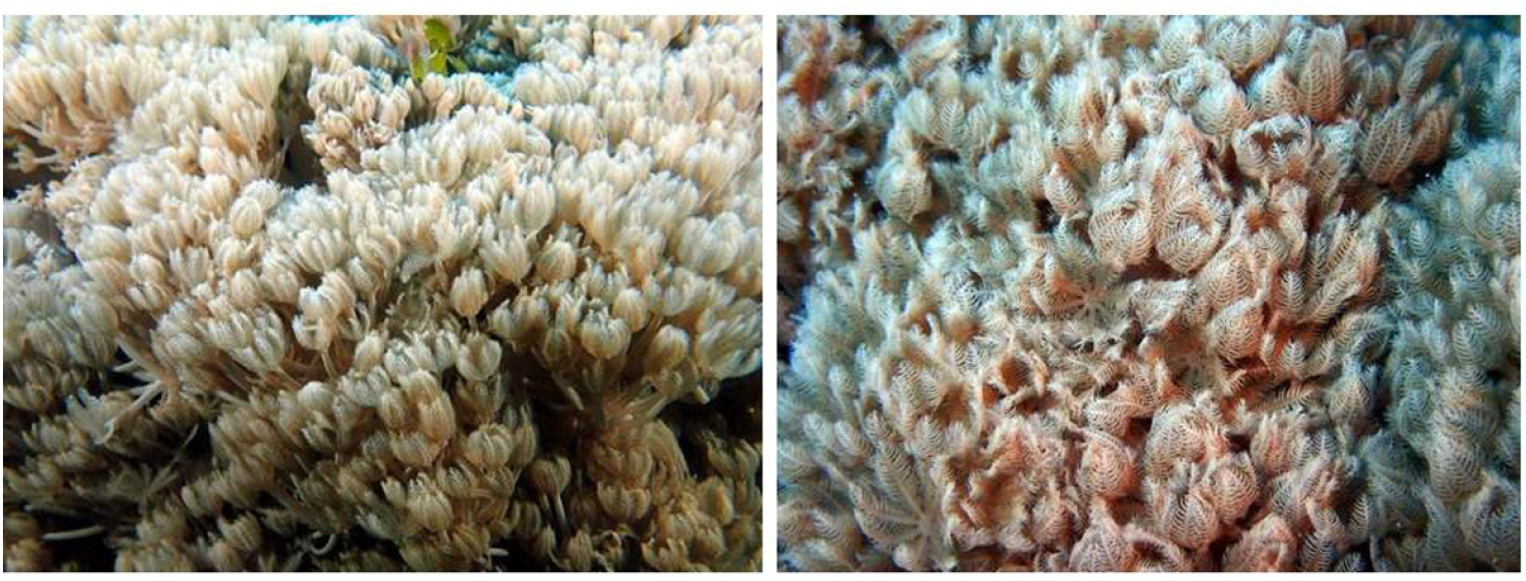
Details of polyps of *Xenia umbellata* in Boca de Calderas, Mayabeque, October 5, 2023. Photo credit: R. Espinosa Sáez, J. Estrada Estrada, A. Martínez Pérez, Y. del Carmen Luguera González.

Espinosa *et al*. (2023) tentatively identified the invasive octocorals as *Unomia stolonifera* based on morphological characteristics; however, they emphasized that definitive identification would require DNA analyses from the colonies. In parallel, two new allochthonous octocoral species (family Xeniidae) were reported in Puerto Rico (Toledo-Rodriguez *et al*. 2025 a, b), and communication and collaboration among stakeholders were established to discuss species identification and appropriate eradication strategies to be implemented. Correct taxonomic identification was considered paramount for effective invasive species management, as it provides the basis for selecting appropriate response actions, offers insight into potential introduction pathways or modes of propagation, and reduces the risk of ineffective or misdirected interventions that may further compromise the coastal ecosystem. However, identifying xeniid species can be complicated when based solely on external and internal morphology. Overlapping shapes of colony forms, stalk growth patterns, and characteristics of the pinnate tentacles make them difficult to distinguish visually, especially in the field. Unlike most other octocoral taxa that can be identified from the characteristic shapes and distribution of sclerites (calcitic skeletal elements), the sclerites of xeniids are very homogeneous and exhibit few species-specific characters that can be observed under light microscopy (Halasz *et al*. 2019). These limitations emphasize the importance of integrative taxonomy, which requires molecular analyses. In this study, we are fulfilling the need for molecular data employing multi-loci barcoding to clarify the taxonomic identity of the Cuban specimens.

## Methods

### Sampling Sites

Samples were collected during surveys conducted by personnel of the Institute of Marine Sciences (ICIMAR), the Antonio Núñez Jiménez Foundation for Nature and Humanity (FANJ), the Center for Marine Research (CIM-UH) and the Rincón de Guanabo Protected Natural Landscape Protected Area (PNPRdG). The analyzed samples were collected at Boca de Calderas, Province Mayabeque, near PNPRdG Rincón de Guanabo (82º14’39,7’’ W; 23º10’47,4’’ N). Samples were sent to personnel of the Puerto Rico Department of Natural and Environmental Resources, who then distributed them to the McFadden Laboratory, Harvey Mudd College and the Marine Genomic Biodiversity Lab at the University of Puerto Rico, Mayagüez for molecular identification.

### DNA procedures and analysis

For each polyp, 4-5 tentacles were excised using sterile tweezers and a scalpel, then used as a tissue source for DNA extraction. DNA extraction was performed using the Qiagen DNeasy Blood and Tissue kit with manufacturer’s protocol, or Qiagen DNeasy PowerSoil® kit with an additional homogenization step consisting of three sets of 20 seconds, with 1-minute intervals on ice between the sets to maximize DNA yield. PCR amplifications of three mitochondrial regions (16S/ND2, mtMutS, COI) and the nuclear large ribosomal subunit (28S rRNA) were done using the primers listed in Koido *et al*. (2022). For the PCR amplifications, we followed the protocol in Toledo-Rodriguez *et al*. (2025a). Amplified DNA was sequenced in both directions in an Applied Biosystems SeqStudio Genetic Analyzer using the Big Dye 3.1 Terminator Cycle chemistry. The DNA traces were inspected for quality and accuracy in Codon Code Aligner v. 10.0.2 (Codon Code Corp.). GenBank Accession Numbers are 16S/ND2 XXXXXXX, mtMutS XXXXXXX, COI XXXXXXX, and 28S sequences XXXXXXXX.

The resulting DNA sequences were concatenated and then aligned in MAFFT v.7 (Katoh & Standley 2013) with default parameters. We compared the DNA sequences from Cuba with 31 published xeniid sequences in a phylogenetic analysis. The analysis included selected 16S/ND2, mtMutS, COI, and 28S sequences from McFadden *et al*. (2017) and Benayahu *et al*. (2022) (Supplementary Table 1). A maximum likelihood (ML) tree was constructed using raxmlGUI 2.0.17 (Stamatakis 2014, Edler *et al*. 2020) and rooted with *Xenia umbellata*, the sister group of *Ovabunda*. First, the program modeltest-ng (Darriba *et al*. 2020) was used in raxmlGUI to estimate the best nucleotide substitution model for the concatenated data. The best nucleotide substitution model (TPM3uf+I) for the alignment was selected using the Akaike Information Criterion. Then, an ML + thorough bootstrap search for the best tree was conducted in raxml-ng (Kozlov *et al*. 2019). Clade support was assessed with 100 bootstrap replicates (Felsenstein 1985). The consensus tree was viewed and edited in FigTree 1.4.4, then imported into Adobe Illustrator 30.2.1 to improve visualization.

## Results

Sequences obtained for each of four loci used for DNA barcoding of octocorals (16S/ND2 = 710 bp; mtMutS = 714 bp; COI = 771 bp; 28S rDNA = 708 bp, a total of 2,903 bp) were 100% identical to barcodes published previously for *Xenia umbellata* from the Red Sea and from the invasive population in Puerto Rico (Fig. 6). Photographs of the Cuban colonies taken in situ (Figs. 3, 4, 5) further support this identification, showing all polyps localized to the tops of the colony stalks, a diagnostic character of the genus *Xenia*. In contrast, *Unomia stolonifera* exhibits isolated polyps on the stalks and base of the colony (Benayahu *et al*. 2021).

**Figure 6.**
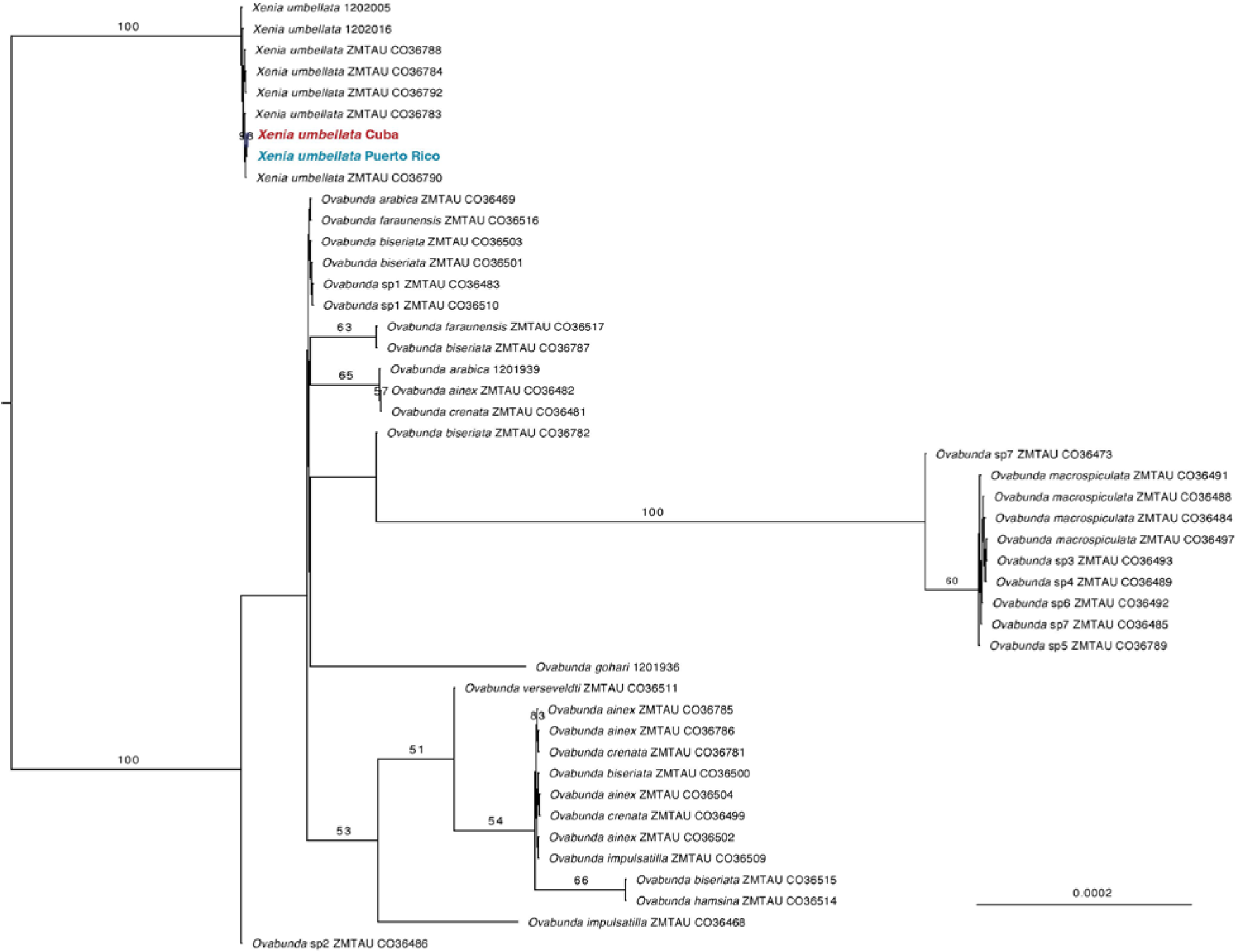
A maximum likelihood tree based on concatenated sequences of mtMutS, 16S/ND2, COI, and 28S gene regions showing the phylogenetic placement of invasive xeniid corals in Cuba (in bold red) and in Puerto Rico (in bold blue) within the *Xenia umbellata* clade. The tree was rooted on the *Xenia*-*Ovabunda* branch. Numbers on clades represent parametric bootstrap values (Bp > 50). Voucher information follows the specimen’s name (see McFadden *et al*. 2017 and Supplementary Table 1).

## Discussion

The occurrence of invasive octocorals from the family Xeniidae is evident in the wider Caribbean and causes ever increasing concerns in the region (Ruiz-Allais *et al*. 2014, 2021; Espinosa *et al*. 2023; Toledo-Rodriguez *et al*. 2025a, b; Veglia *et al*. 2025). Since the detection of xeniid colonies in Cuban waters, initially identified morphologically as *Unomia stolonifera*, an immediate response was executed to limit further spread and potential ecological impact. This response was motivated by the ecological impacts reported in Venezuela, where *U. stolonifera* rapidly expanded and monopolized benthic habitats (Ruiz-Allais *et al*. 2014, 2021). More recently, in Puerto Rico, similar concerns emerged regarding invasive octocoral colonies initially presumed to be *U. stolonifera* but later confirmed by molecular analysis to be two separate species of pulsing corals, *X. umbellata* (Toledo-Rodriguez *et al*. 2025a; Veglia *et al*. 2025) and *Latissimia ningalooensis* (Toledo-Rodriguez *et al*. 2025b). Here, using identical approaches applied to Puerto Rico specimens, we examined four DNA regions to confirm unequivocally that the invasive Cuban species is not *U. stolonifera* as originally reported (Espinosa *et al*. 2023), but identified instead as *X. umbellata*.

The phylogenetic footprint of *X. umbellata* is unique within Xeniidae (McFadden *et al*. 2017) and standard DNA barcode sequencing can provide definitive evidence for the presence of the species in the area (McFadden *et al*., 2019). Our results therefore confidently indicate that *X. umbellata* may be more widespread in the region than previously thought, with additional populations possibly occurring along the coasts of the Dominican Republic and Haiti, or in additional locations in the Caribbean. This discovery of *X. umbellata* in Cuba and Puerto Rico has changed the prevailing narrative regarding soft coral invasion in the Caribbean. Rather than addressing a single species scenario—*U. stolonifera*, which has rapidly expanded from Venezuela (Ruiz-Allais *et al*., 2014; 2021) to the southern Caribbean—the region is now confronted with the introduction of multiple species: *Xenia umbellata* (Toledo-Rodriguez *et al*. 2025a; Veglia *et al*. 2025) and *Latissimia ningalooensis* (Toledo-Rodriguez *et al*. 2025b) in Puerto Rico, and *X. umbellata* in Cuba (this study). This represents a conservative scenario, as *U. stolonifera* is likely to continue its range expansion across the Caribbean, beyond Venezuela.

The first octocoral aggregations (presumably also *X. umbellata* since we unequivocally confirmed the identification a colony only from Boca de Calderas) were discovered in Bacuranao Inlet, east of Havana, in September 2022, nearly a year before the first colonies were observed by recreational divers in southern Puerto Rico in October 2023. However, based on the size of the aggregations, at least in Puerto Rico, the initial sightings likely postdate the actual establishment of colonies. The near simultaneous detection of *X. umbellata* in two Caribbean islands does not point to secondary spread from one place to another. Together, the available records may be consistent with independent introductions rather than with direct spread between Cuba and Puerto Rico. Two lines of evidence support this interpretation. First, traffic of commercial vessels between the two islands is very limited or nonexistent, severely reducing the possibility of spreading between the sites via shipping. Rather, a scenario involving independent introductions associated with maritime infrastructure (e.g., in ballast water or on fouled hulls or fishing gear) appears somewhat plausible. This is supported by the proximity of the two Cuban soft coral sites (Fig. 1) to the port of Havana: approximately 12 km from Bacuranao Inlet and 27 km from Boca de Calderas, both located east of the port. Similarly, the first documented patches of *X. umbellata* in Puerto Rico were discovered near the port of Ponce, a busy hub of international maritime traffic (Toledo-Rodriguez *et al*. 2025a). Second, based on near-surface buoy drifter data, there is a persistent westward surface flow across the Venezuelan Basin extending as far as the Jamaican Ridge, followed by a northwestward flow from there to the Yucatán Channel (Centurioni & Niiler 2003). This circulation pattern suggests that propagules would be transported from east to west—that is, from Puerto Rico toward Cuba—rather than in the opposite direction. Though the invasion sites in Puerto Rico are located on the south side of the island so far, whereas in Cuba they were encountered on the north side, suggesting that, if the east-to-west propagule scenario is possible, there may be additional soft coral aggregations on the southern coast of Cuba. None have been reported yet from the southern coast, even though between 2023 and 2025 several reef surveys covered the entire southern part of Cuba, under the auspices of “Bojeo a Cuba 2023”, a scientific expedition of the Cuban archipelago. It is noteworthy that the known distribution of the invasive octocorals in both jurisdictions is limited to occasional reports from recreational divers or researchers targeting specific areas, rather than through a standardized monitoring effort, and is thus likely underestimated. Further, misidentification with native organisms is another factor that may contribute to the lack of reporting in both Cuba and Puerto Rico.

Although the precise route of *X. umbellata* introduction into Cuba remains uncertain, several other pathways are plausible, with the marine aquarium trade considered the most likely hypothesis above all, as has been suggested for other invasive xeniid octocorals (Ruiz-Allais *et al*. 2014; Mantellato *et al*. 2018; Menezes *et al*. 2022; Bolick and Lee 2023). Reports from 2025, although not yet verified in the field, indicate the presence of three new populations intentionally planted by aquarists. Other possible pathways of introduction cannot be ignored including biofouling, ballast water transport, floating debris, macroplastics, *Sargassum* (e.g. Bailey 2015; Mantelatto *et al*. 2018; García-Gómez *et al*. 2021; Menezes *et al*. 2022; Hoeksema *et al*. 2023).

Our study does not rule out the presence of *U. stolonifera* in Cuba; rather, it provides evidence that a sample from the second documented octocoral aggregate in Boca de Calderas—initially thought to be *U. stolonifera*—was unequivocally identified as *X. umbellata* through molecular barcoding. The sample was randomly collected by divers as representative of the observed octocoral aggregations. Therefore, unless new data indicate otherwise, we assume that all other sightings in the area correspond to *X. umbellata*. Nevertheless, marine vessel traffic between Venezuela and Cuba continued uninterrupted through at least mid-2020, providing a potential pathway for accidental transport of *U. stolonifera* propagules and other species via ballast water or on contaminated gear (Ruiz-Allais *et al*. 2021). Therefore, marine resource managers in Cuba should remain vigilant for the possible introduction of *U. stolonifera* or other marine invaders into the country’s coastal ecosystems.

Early detection and aggressive intervention efforts may still reduce the expansion of these invasive xeniids. Eradication efforts in Cuba have been successful in managing recurrences at a particular site through contact surveillance and action (Espinosa *et al*. 2023); however, given the regenerative capabilities of *X. umbellata* (Nadir *et al*. 2023), the action must continue. Reports for other areas of Cuba could not be confirmed due to limited resources. In Puerto Rico, the distribution is confirmed to be widespread along the southwest coast, with large colony sizes (unpub. data). Coordinated eradication was implemented but was limited due to a lack of funds. As expected, the areas were repopulated by presumably untreated nearby aggregations (unpub. data). Xeniids are spreading across the Caribbean. With adequate funding, perseverance, and collaboration, these soft corals could be eradicated—or at least controlled—if early detection is achieved. A coordinated monitoring approach that integrates wide-area autonomous underwater vehicle (AUV) surveys, remotely operated vehicles (ROVs), environmental DNA (eDNA) sampling, and diver-based ground-truthing through direct observations and specimen collections would establish its current distribution and early detection of outbreaks in areas where it has not been reported, helping safeguard Caribbean coastal ecosystems.

## Acknowledgments

We thank Dr. Silvia Patricia González Díaz (University of Havana) for reviewing an earlier version of this manuscript. Valerie Miller (Environmental Defense Fund) assisted in critical communication among the authors. This publication was made possible with support from the Sequencing and Genomics Facility of the UPR Río Piedras & MSRC/UPR, funded by NIH/NIGMS-Award Number P20GM103475.

